# Chronic intermittent alcohol yields sex-specific disruptions in cortical-striatal-limbic oscillations

**DOI:** 10.1101/2024.08.23.609453

**Authors:** Kelly A. Hewitt, Skylar E. Nicholson, Madilyn J. Peterson, Lucas L. Dwiel, Angela M. Henricks

## Abstract

**Background:** While the neurobiology of alcohol use disorder (AUD) has been extensively researched, the vast majority of these studies included only male organisms. However, there are significant sex differences in both the causes and consequences of alcohol misuse and dependence, suggesting sex-specific neurobiological mechanisms. The current study used a rodent model to determine whether chronic alcohol exposure impacts sex-specific neural circuits, and whether these changes contribute to the development of alcohol misuse.

**Methods:** Male and female Sprague-Dawley rats were trained to self-administer 10% alcohol before implanting bilateral electrodes into the infralimbic medial prefrontal cortex (IL), nucleus accumbens shell (NAcSh), and central nucleus of the amygdala (CeA). Half of the rats were then exposed to four weeks of chronic intermittent alcohol (CIA) vapor (14 hours on/10 hours off). During acute withdrawal (6-8 after the vapor turns off), local field potentials (LFPs) were recorded from the IL, NAcSh, and CeA during 30-minute self-administration sessions. Using an unbiased machine learning approach, we built predictive models to determine whether/which LFP features could distinguish CIA-exposed from control rats in each sex, as well as if any of these LFP features correlated with rates of alcohol self-administration.

**Results:** Female rats self-administered more alcohol in general compared to males, but only males exposed to CIA showed increased alcohol intake during acute withdrawal. LFPs predicted CIA exposure in both sexes better than chance estimates, but models built on IL and NAcSh oscillations performed the best in males, while models built on IL and CeA LFPs performed best in females. High γ LFPs recorded in the NAcSh correlated with rates of alcohol self-administration in males exposed to CIA, while only left-right NAcSh β coherence correlated with drinking in control females.

**Conclusions:** These data provide support for the hypothesis that the neural circuits driving alcohol dependence development are sex-specific, and that high frequency oscillations in the NAcSh may be related to the increased drinking observed in males exposed to CIA. Overall, these data add to our understanding of the neurobiological underpinnings behind the sex differences observed in AUD and offer promising biomarkers for future therapeutic research.

## BACKGROUND

Alcohol use disorder (AUD) is the most prevalent substance use disorder in the United States, causing serious social, behavioral, and health-related problems (1,2). Despite several FDA-approved treatments for AUD (3), relapse is still extremely common with estimated long-term relapse rates as high as 85% (4).

There are significant sex differences in the reasons for drinking and the physical, behavioral, and neurobiological consequences of chronic alcohol misuse. While men consume more alcohol and are more likely to be diagnosed with AUD compared to women, this gender gap has begun to shrink over the last decade, with more women engaging in risky alcohol drinking (5–7). Commonly used medications for AUD, which were developed and tested primarily in male animals, are less effective in women compared to men (8,9). In fact, the vast majority of preclinical neuroscience research aimed at understanding AUD has not included female animals, which may contribute to poor translatability from preclinical animal research to clinical treatment development (10). Therefore, there is a vital need for a more inclusive and thorough understanding of the neurobiological mechanisms driving alcohol misuse in both sexes.

We have previously demonstrated that local field potentials (LFPs) collected in the nucleus accumbens shell (NAcSh) and medial prefrontal cortex (mPFC) predict the volume of alcohol consumed in non-dependent male, but not female, rats (11). LFPs reflect the dynamic flow of information across neural circuits, capturing electrical information within (power) and between (coherence) brain regions that provides insight into how synchronized/asynchronized activity is across areas of interest (12,13). LFPs can be collected in awake, freely behaving animals over long periods of time, and therefore have the potential to serve as biomarkers for different behavioral states (14). Our previous results suggest that cortical-striatal oscillations may be particularly relevant to male, but not female, drinking behavior. While the mPFC and NAcSh play vital roles in voluntary alcohol consumption, contributing to the rewarding, motivational, and cognitive components of AUD (15), other circuits play important roles as well and may be more relevant to females.

Women are more likely to use alcohol to alleviate stress (negative reinforcement), while men are more likely to use alcohol as a positive reinforcer (16). Further, in both humans and rodents, females are more susceptible to stress-induced relapse (17). We therefore hypothesize that oscillations recorded in stress-related brain regions, like the amygdala, may contain more information regarding drinking behavior in females.

In order to collect data relevant to AUD, the current study employed a commonly used model of alcohol dependence in rodents: chronic intermittent alcohol vapor exposure (CIA). The main objective was to determine whether CIA alters LFPs recorded in the infralimbic (IL) mPFC, NAcSh, and central nucleus of the amygdala (CeA) in a sex-specific manner. Using an unbiased machine-learning approach, we hypothesized that IL, NAcSh, and CeA LFPs would predict whether animals were exposed to CIA and correlate with rates of alcohol self-administration, with cortical-striatal and cortical-limbic LFPs containing the most information for males and females, respectively.

## METHODS

### Animals

Adult (approximately 70 days old) male and female Sprague-Dawley rats were used for these experiments (n = 6-7/group/sex; 27 total). Animals were obtained from Inotiv (West Lafayette, IN) and were housed in standard polycarbonate cages with *ad libitum* access to food and water. Rats were kept under a 12-hour reverse light-dark cycle (lights off at 08:00), allowing for all experimental procedures to be conducted during the rat’s active phase and to avoid potential circadian rhythm disruptions. Animal facilities were accredited by the American Association of Laboratory Animal Care (AALAC), and all procedures were approved by the Institutional Animal Care and Use Committee (IACUC) at Washington State University. Animals were kept in same-sex pairs prior to electrode implantation and single housed after surgery to prevent damage to the electrodes. All animals were handled daily for a minimum of 5 days the week prior to the start of testing.

### Alcohol self-administration training

Self-administration took place in standard, sound attenuating operant boxes (10” L x 12.5” W x 8.5” H) programmed using MED-Associates IV software. Boxes are equipped with two retractable levers on the same wall of the chamber with a center receptacle for liquid distribution. Presses to the active lever resulted in the delivery of 0.1mL of liquid. Responses on the inactive lever resulted in no reinforcer. A house light was illuminated for the duration that levers were available for each session.

Rats were trained to self-administer 10% ethanol in water under a fixed ratio-1 (FR-1) reinforcement schedule using a sucrose fade technique, as described in our previous work (18,19). Rats were trained with the following sequence of solutions: 10% sucrose, 10% sucrose + 10% ethanol, 5% sucrose + 10% ethanol, and finally 10% ethanol. Each session in the operant chambers lasted 30 minutes, and sessions were conducted 5 days per week, Monday-Friday. Training was considered complete after 15 sessions of 10% ethanol self-administration. The average g/kg of alcohol consumed over the last four sessions was calculated for each animal as the baseline drinking level.

### Electrode implants

Following self-administration training, rats were anesthetized with isoflurane gas before electrodes were implanted via stereotaxic surgery. Electrodes were constructed in-house, and designed to collect LFPs from the NAcSh (from bregma: DV -8.0mm; AP +1.2 mm; ML +/- 1.0mm) and IL mPFC (from bregma: DV –4.5mm, AP +3.4mm, ML +/-0.75mm), as outlined previously (11,20), as well as the CeA (from bregma: DV –8.5mm, AP –2.0mm, ML +/-3.5mm). Four stainless steel screws were attached to the skull around the electrode site and dental acrylic (Lang Dental, Wheeling, IL, USA) was used to secure the electrode in place. No rats had to be excluded based on electrode placement (see *Histology*). Animals were allowed to recover for at least seven days before the next phase of the experiment.

### Chronic intermittent alcohol exposure

After recovery, animals were either exposed to CIA vapor (n = 6-7/sex) or normal room air (control; n = 7/sex) for four weeks, as described previously (18,19). Our custom-built, 4-chamber apparatus (La Jolla Alcohol Research Inc., La Jolla, CA, USA) allows blood alcohol levels (BALs) to be titrated by the experimenter by adjusting the rate of 95% ethanol that is vaporized and integrated into the air flow supplying the sealed Plexiglas chambers in which the rats were housed (25.5” L x 21” W x 13” H). Food and water were available *ad libitum*. CIA animals were exposed to vaporized ethanol for 14 hours per day from approximately 18:00-08:00; for 10 hours a day they were just exposed to room air that was filtered into the chamber. Animals in the CIA condition remained in the vapor chambers for the entirety of the experiment. Blood was collected at least two times per week to ensure BALs remained at an appropriate level (between 150-250mg%), as described previously (18,19). BALs were determined by centrifuging approximately 0.1mL of blood collected from the tail vein. Plasma samples were then analyzed for ethanol content using the Analox AM1 (Analox Instruments Ltd., Lunenburg, MA, USA).

### Alcohol self-administration after CIA

Following the four-week period of initial CIA exposure, all rats completed four baseline alcohol self-administration sessions (with a minimum of 1 day between sessions) during acute alcohol withdrawal (∼7 hours after the vapor turned off).

### Collecting local field potentials

Next, three separate alcohol self-administration sessions were completed while collecting LFP data with an OmniPlex Neural Recording Data Acquisition System (Plexon, Dallas, TX, USA). Time-locked behavior was also recorded via a CinePlex camera system (Plexon, Dallas, TX, USA). Twelve rats completed an additional session due to technical issues during one of the original recording sessions. Signals passed through a digital head stage and custom-made cable connected to a commutator, which allowed free movement and behavior while tethered to the data acquisition system in the operant chamber.

### Histology

At the end of the experiment, rats were euthanized using CO_2_ gas and brains were snap frozen in 2-methylbutane on dry ice. Tissue was stored at -20℃ prior to sectioning. Tissue sections were then stained with thionine and electrode placement was verified using a Leica CTR 6500 microscope (Wetzler, Germany) (Figure 1).

**Figure 1:**
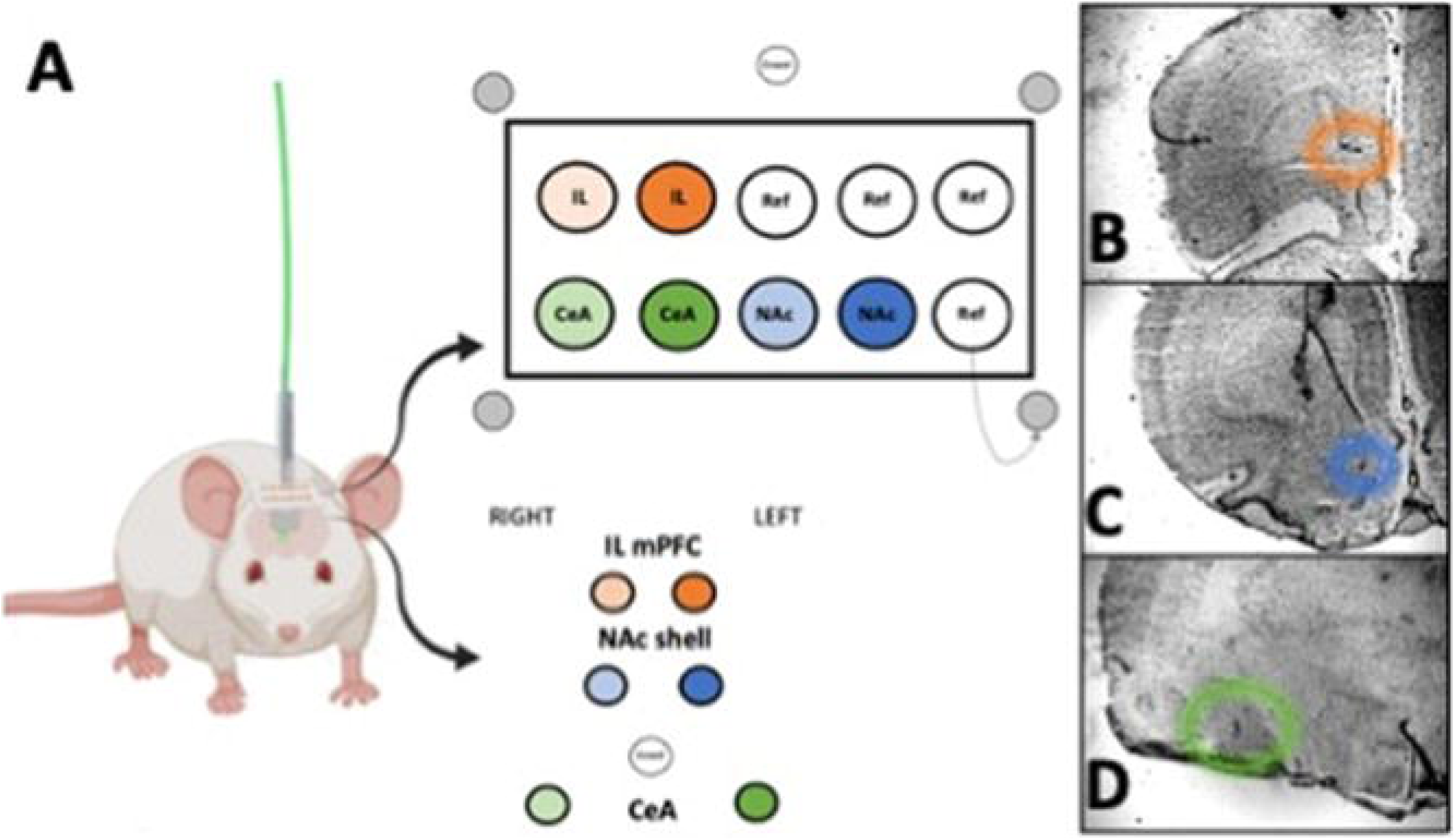
Electrode schematic (A) and example placements in the IL (B), NAcSh (C) and CeA (D).

**Figure 2:**
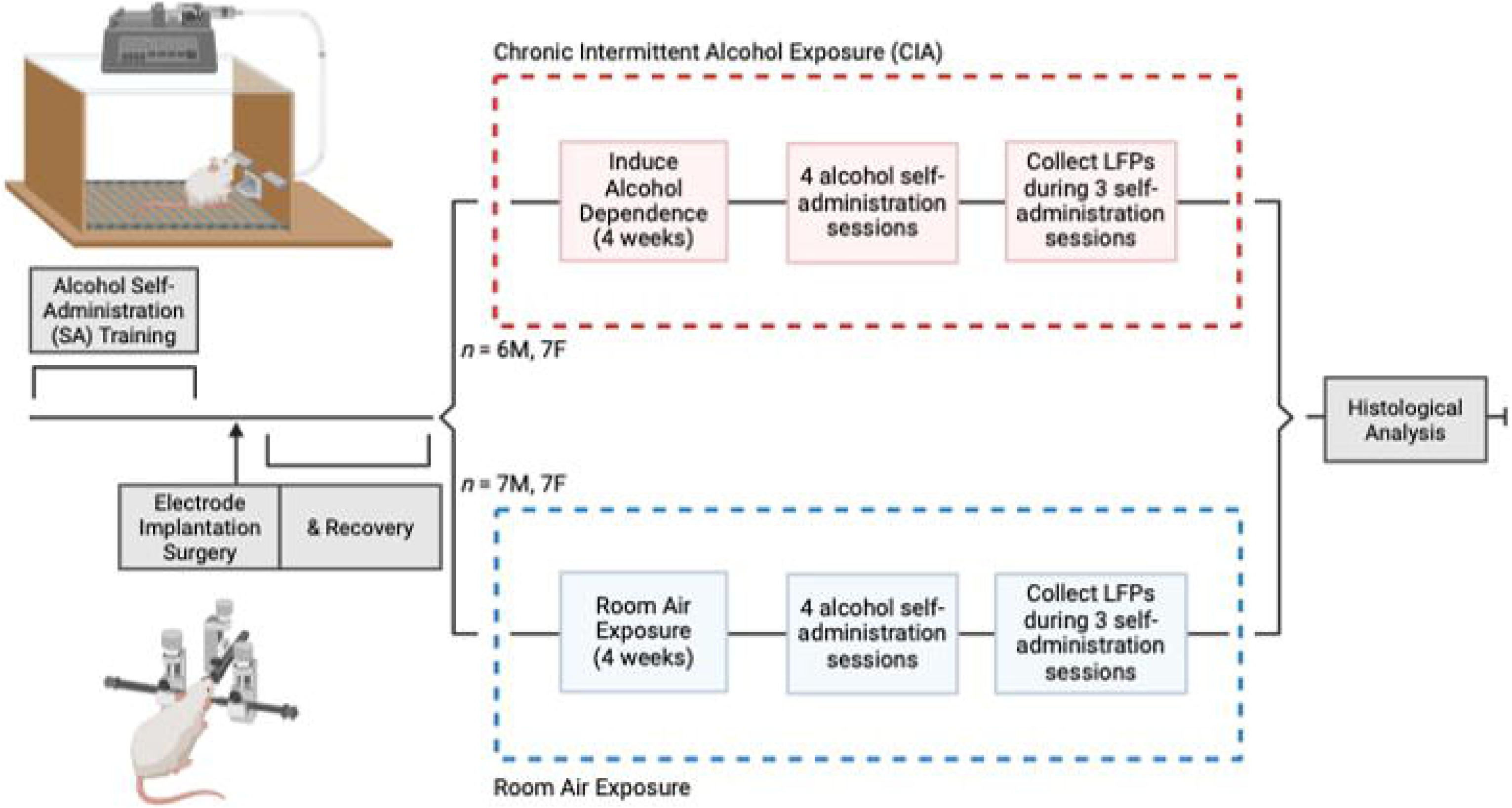
Experimental timeline.

### Statistical Analyses

#### Alcohol self-administration

To determine how alcohol self-administration was impacted by CIA, a 2 x 5 mixed factorial ANOVA was conducted with dependence state (controls, CIA) as the between-subjects factor, and session as the within-subjects factor (pre-CIA baseline, plus 4 post-CIA sessions), and g/kg of alcohol consumed as the dependent variable. To see how animals behaved during LFP collection, a 2 x 3 mixed factorial ANOVA was conducted with dependence state (controls, CIA) as the between-subjects factor, three LFP self-administration sessions as the within-subject factors, and g/kg of alcohol consumed as the dependent variable. Because we had an *a priori* hypothesis that there would be sex differences in behavior, based on our previous data (18), we performed analyses for each sex separately.

#### Unbiased prediction modeling

Custom code written in MATLAB R2023a allowed for the characterization of the LFP power spectral densities (PSDs) within and coherence between brain regions for each rat and each session, as previously reported (11,20,21). Data from each recording was analyzed using frequency ranges from the rodent literature (δ =1-4 Hz, θ = 5-10 Hz, α = 11-14 Hz, β = 15-30 Hz, low γ = 45-65 Hz, and high γ = 70-90 Hz) (22,23). After applying a 60Hz noise-filter, PSDs and coherence for each 5-second segment were averaged together across each 30-minute recording session. We used a penalized regression model (lasso) to predict binary (CIA vs. controls) and continuous (g/kg of alcohol consumed) outcomes using IL, NAcSh, and CeA LFPs, as outlined previously (20).

Each recording session produced 126 LFP features: 36 measures of power (6 frequency bands x 6 channels) and 90 measures of coherence (6 frequency bands x 15 channel combinations). The MATLAB package Glmnet was used to implement the lasso using a 5-fold cross-validation. Models were trained on 80% of the data and tested on a left-out 20%, which was then repeated 100 times with different, randomly selected sets of training and testing data. The real models were then compared to 100 random permutations of the data (permuted models), as described previously (21). Real and permuted models were generated for each sex separately. Additionally, we performed logistic regressions on each LFP feature individually to determine the relative predictive accuracy of said feature, as we have previously described in detail (21). We further used simple linear regressions to determine whether each of the top five LFP features with the highest prediction accuracy correlated with g/kg of alcohol consumed in self-administration sessions for each group (male control, male CIA, female control, female CIA). Lastly, we used moderation analyses to determine if CIA exposure influenced the strength and directions of any correlations.

## RESULTS

### Alcohol self-administration

For males, a mixed factorial ANOVA revealed a significant effect of session [*F*(4,44) = 4.01, *p*<0.01, 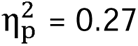], a significant effect of group [*F*(1,11) = 7.68, *p*<0.05, 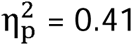], and a significant session*group interaction [*F*(4,44) = 3.27, *p*<0.05, 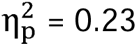] for g/kg of alcohol consumed. Post-hoc tests showed that CIA-exposed males drank more alcohol in sessions 3 and 4 (*p*<0.05; Figure 3A) compared to control males.

**Figure 3:**
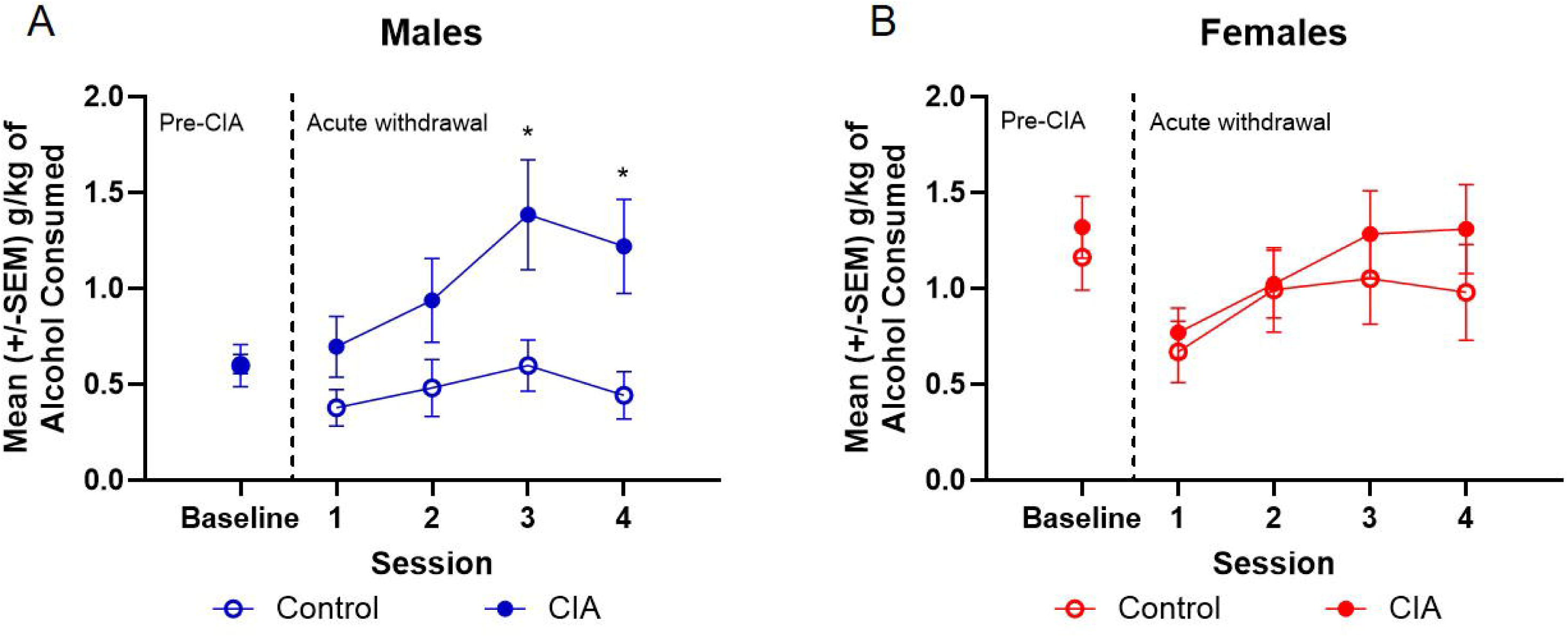
Average g/kg of alcohol consumed in male (A) and female (B) rats exposed to CIA or room air (control) during 30-minute self-administration sessions before (pre-CIA baseline) and after (acute withdrawal) four weeks of CIA exposure. CIA led to increased drinking in the last two self-administration sessions in males (\**p*<0.05; n=6-7/group), but no changes in female drinking (*p*>.05; n=7/group).

For females, a mixed factorial ANOVA revealed a significant effect of session [*F*(4,48) = 3.75, *p*<0.05, 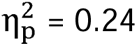], but no effect of group [*F*(1,12) = 0.65, *p*=0.44, 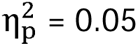] or a group*session interaction [*F*(4,48) = 0.30, *p*=0.88, 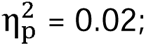 Figure 3B] for g/kg of alcohol consumed. These data are consistent with our and others’ previous findings (18,26). It is also important to note that, in both sexes, inactive lever presses did not differ by group (Supplemental Figure 1).

During LFP collection, tethering the animals to the commutator eliminated any session or group effects on g/kg of alcohol consumed, and drinking was suppressed below baseline levels (Supplementary Figure 2).

### Unbiased prediction models

Using LFPs from all regions (IL, NAcSh, and CeA), Figure 4 shows the average area under the receiver operating characteristic curve (AUC) for the models predicting CIA vs. control exposure in males (real AUC = 0.63 ± 0.04, permuted AUC = 0.52 ± 0.04; 4A) and females (real AUC = 0.81 ± 0.03, permuted AUC = 0.50 ± 0.03; 4B). We also determined the prediction accuracy of models built on just cortical-striatal vs. just cortical-limbic LFPs. In males, models built on IL and NAcSh LFPs (real AUC = 0.78 ± 0.04, permuted AUC = 0.49 ± 0.04) performed better than models built on IL and CeA LFPs (real AUC = 0.65 ± 0.04, permuted AUC = 0.48 ± 0.03; Figure 4A). The opposite was true in females; models built on IL and CeA LFPs (real AUC = 0.83 ± 0.03, permuted AUC = 0.49 ± 0.03) performed better than models built on IL and NAcSh LFPs (real AUC = 0.57 ± 0.04, permuted AUC = 0.49 ± 0.03; Figure 4B).

**Figure 4:**
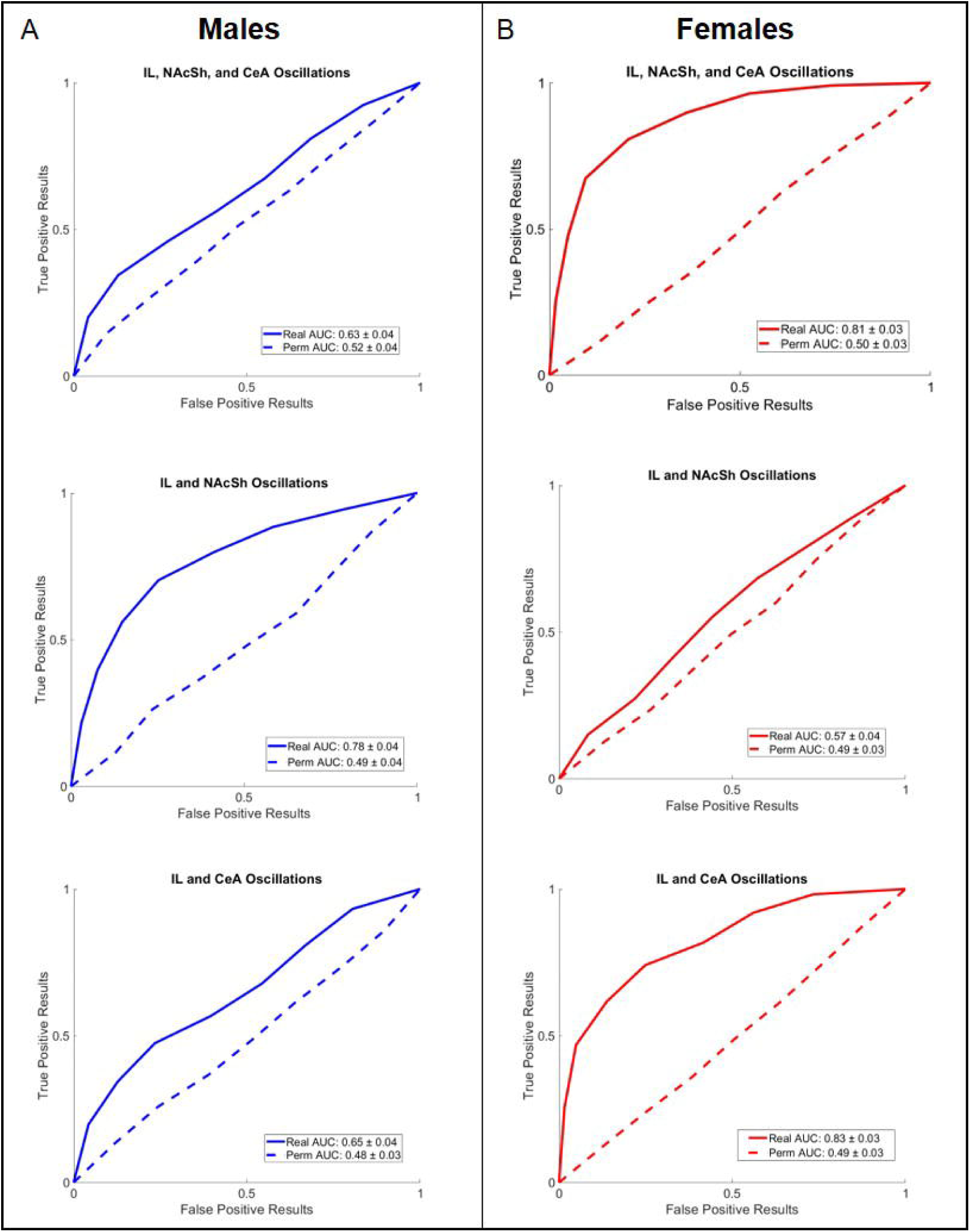
Average area under the receiver operating characteristic curve (AUC) +/- SD for real and permuted binary prediction models (n = 6-7/sex × 3 recordings/rat) in males (A) and females (B). Models built from IL and NAcSh LFPs best predicted whether males were exposed to CIA. Models built from IL and CeA LFPs best predicted whether females were exposed to CIA.

Table 1 indicates the top five LFP features with the highest prediction accuracies distinguishing CIA vs. control exposure for each sex, including the AUCs for each feature. IL and NAcSh LFPs best predicted CIA exposure in males, while primarily CeA LFPs best predicted CIA exposure in females.

**Table 1.** Neural features that best predict CIA exposure.

| Males |  | Females |  |
| --- | --- | --- | --- |
| Feature | Mean AUC | Feature | Mean AUC |
| $IL_R - NAcSh_L \text{ } l\gamma$ | 0.75 | $CeA_L \text{ } \alpha$ | 0.87 |
| $NAcSh_R \text{ } h\gamma$ | 0.75 | $CeA_R \text{ } \alpha$ | 0.81 |
| $IL_R \text{ } \delta$ | 0.74 | $CeA_L \text{ } h\gamma$ | 0.80 |
| $IL_L - CeA_L \text{ } \theta$ | 0.73 | $NAcSh_L - NAcSh_R \text{ } \beta$ | 0.79 |
| $IL_L \text{ } \delta$ | 0.72 | $CeA_R \text{ } \delta$ | 0.76 |

The top 5 LFP features used in binary models predicting CIA exposure in males and females. Frequency bands [delta ( $\delta$ ), theta ( $\theta$ ), alpha ( $\alpha$ ), beta ( $\beta$ ), low gamma ( $l\gamma$ ), and high gamma ( $h\gamma$ )] are described for power features within and coherence features between neural sites. Subscript indicates left (L) or right (R) hemisphere.

We also attempted to predict g/kg of alcohol consumed within each tethered session from the corresponding LFPs. These prediction models did not outperform chance estimates, likely due to the very low drinking levels exhibited by both sexes when tethered (Supplemental Figure 3).

As an alternative, we measured whether the average value of each of the top neural features correlated with the average g/kg of alcohol consumed during acute withdrawal but prior to tethering (Figure 5). Table 2 shows the correlation coefficients for each of the top neural features in males and females. In males, moderation analyses revealed a significant interaction between right NAcSh high γ power and amount of alcohol consumed (*p*<0.05). Further probing of the interaction revealed that right NAcSh high γ power significantly correlated with g/kg of alcohol consumed in males exposed to CIA [t(9) = -2.89, *p*<0.05], but not in control males [t(9) = 1.05, *p*=0.32] (Figure 5A).

**Figure 5:**
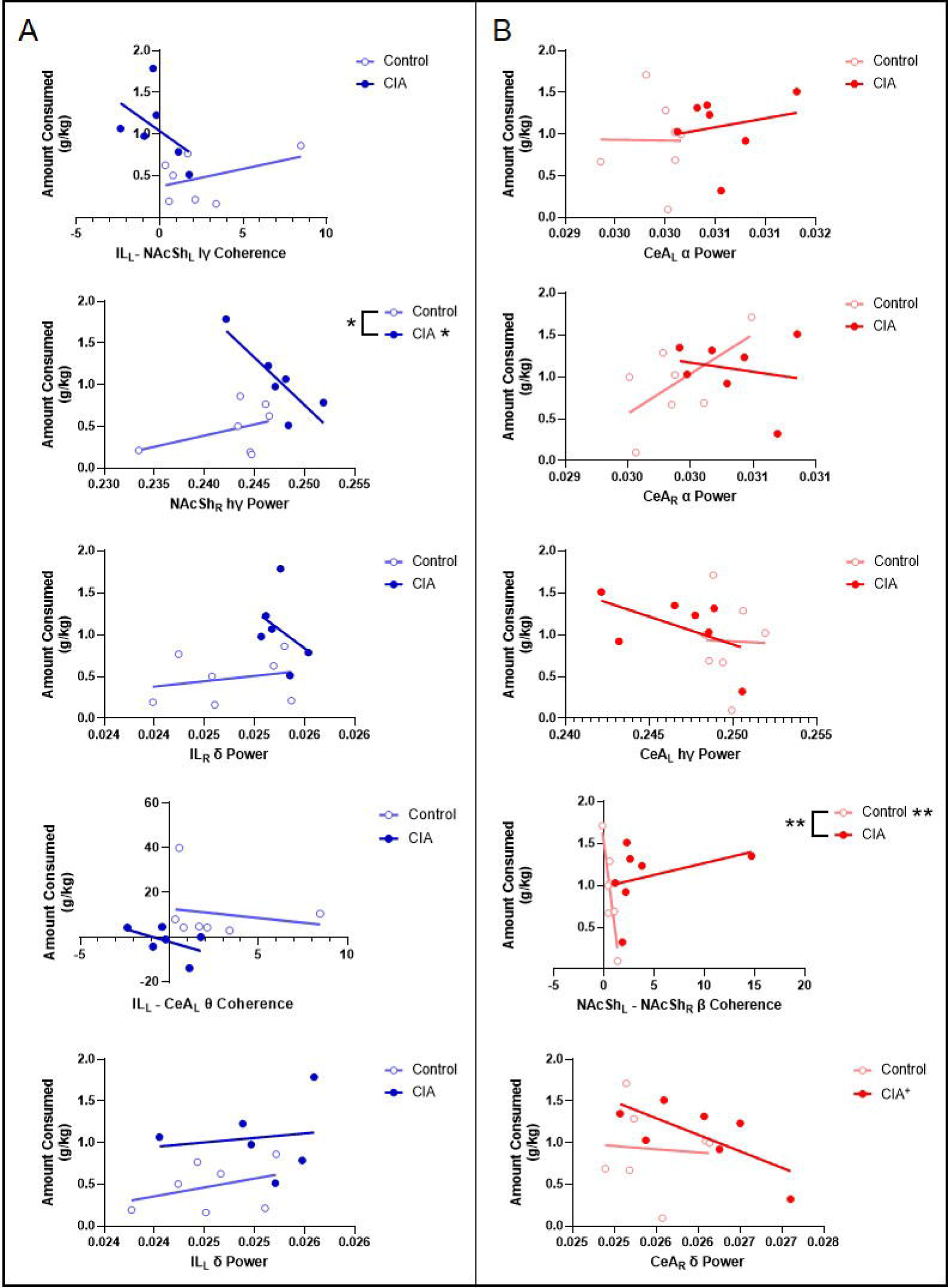
Correlations between average g/kg of alcohol consumed during acute withdrawal in CIA and control rats and average power or coherence across sessions. In males (A), CIA significantly moderated the correlation between alcohol consumption and right NAcSh hγ power, with power and g/kg consumed negatively correlated in CIA males. In females (B), CIA significantly moderated the correlation between alcohol consumption and L-R NAcSh β coherence, with coherence and g/kg consumed negatively correlated in control females. ^+^p<0.08; *p<0.05, **p<0.01.

**Table 2.**
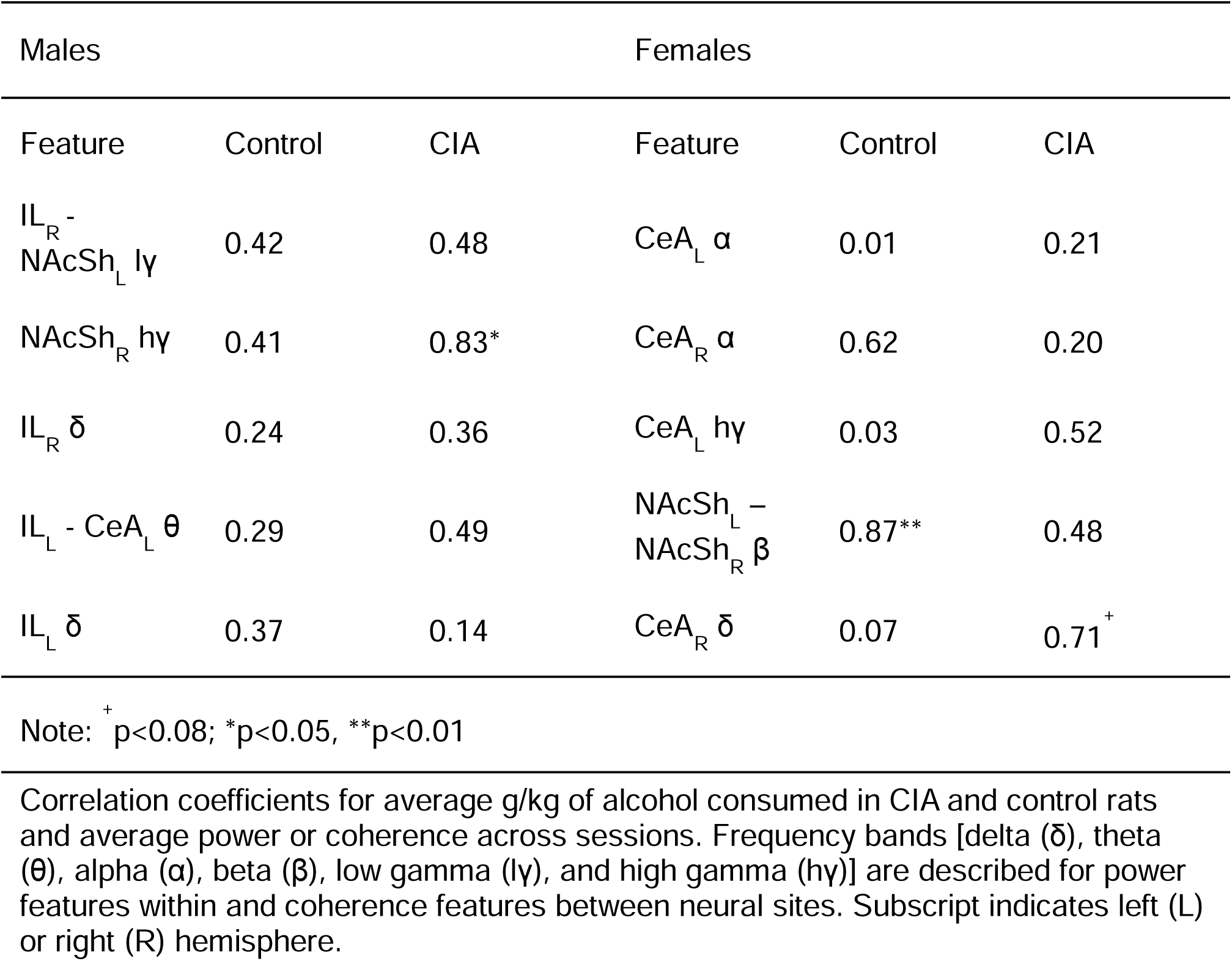
Correlations between average neural feature and g/kg consumed.

In females, moderation analyses revealed a significant interaction between left-right NAcSh β coherence and amount of alcohol consumed in females (*p*<0.01). Further probing of the interaction revealed that left-right NAcSh β coherence significantly correlated with g/kg of alcohol consumed in room air control females [*t*(9) = -3.50, *p*<0.01], but not in females exposed to CIA [*t*(9) = 1.29, *p*=0.23] (Figure 5B). There was also a trend for right CeA δ power to correlate with g/kg consumed in CIA females (*p*=0.08).

## DISCUSSION

Using an unbiased approach, the current study demonstrates that the neural circuits most impacted by CIA are sex-specific. CIA caused disruptions primarily in cortical-striatal regions (IL mPFC and NAcSh) in males and in cortical-limbic regions (IL mPFC and CeA) in females. Further, high γ oscillations recorded in the NAcSh significantly correlated with drinking during acute withdrawal in males exposed to CIA, but not in control males. In females, we identified that β coherence between the right and left NAcSh correlated with drinking in control females, but not in females exposed to CIA. However, there was a trend for a correlation between low frequency (δ) oscillations recorded in the CeA and drinking in females exposed to CIA. It is important to note that though many of our single feature correlations were not significant, our prediction models performed much better than chance estimates, highlighting the power of combining multiple small effects that would usually be undetectable without much larger sample sizes. Overall, these data support the notion that chronic alcohol exposure significantly impacts different neural circuits in males and females, and that high-frequency striatal oscillations may be related to increased alcohol drinking seen in males, which could greatly inform our understanding of the sex differences observed in AUD.

Consistent with our hypothesis, the neural features that contained the most information regarding CIA exposure in males primarily came from LFPs recorded in cortical-striatal regions. This is particularly interesting given the fact that deep brain stimulation (DBS) of the accumbens decreases alcohol consumption in both male rats and men with AUD (20,32,33). Chronic alcohol exposure also leads to dysregulated connectivity between the PFC and accumbens, which may further drive alcohol misuse (34). In line with these findings, CIA exposed males in this study showed lower IL – NAcSh high γ coherence compared to same-sex controls (Figure 5A), suggesting reduced connectivity between those regions. We also showed that high γ power in the right NAcSh significantly correlated with g/kg of alcohol consumed in males exposed to CIA, which is interesting considering that alcohol craving is associated with increased BOLD activity in the accumbens in men, but not in women (35). Further, high frequency oscillations have specifically been tied to substance use; cortical, striatal, and limbic γ power changes correlate with drug delivery or administration of dopamine agonists, suggesting that these oscillations may play a role in motivated behavior and processing reward (22,36–38). Thus, CIA-induced changes to high-frequency cortical-striatal oscillations may be especially relevant to the development of alcohol misuse in males.

On the other hand, the current data suggest that oscillations recorded in the CeA, a region vital to moderating responses to stress and negative affect (39), are significantly disrupted by CIA in females. Increased negative affect plays a critical role in the initiation and maintenance of alcohol misuse (39,40,41). Women also more often report using substances to relieve negative affect compared to men, as well as experience higher levels of subjective stress in general (16,17). In light of the current data, combined with the fact that others have shown that repeated cycles of alcohol withdrawal leads to significant neuroadaptations in the CeA (39,42), we propose that this region is a promising research target for potentially unlocking more information regarding alcohol dependence development in females.

However, although there was a trend for right CeA δ power to correlate with alcohol consumption in CIA-exposed females, none of the CeA-based neural features significantly correlated with drinking. We hypothesize that these findings may be related to inadequate power due to a small sample size in the present experiment, but this requires further investigation.

A novel finding of the current study is that left-right NAcSh β coherence correlated with alcohol consumption in control females. In humans, frontal cortex β oscillations have been associated with genetic variants that predispose an individual to AUD (43). In the NAc, β oscillations are associated with action preparation (44), are inversely related to dopamine availability, and predict relapse to alcohol drinking in rodents (45). While the functional meaning behind NAcSh β coherence is currently unknown, coherence measures are thought to reflect communication between brain regions. We therefore hypothesize that disruptions in inter-accumbens communication may be related to motivation to drink in non-dependent females, but this theory requires significant further investigation.

We also observed significant sex differences in self-administration behavior, which aligns with previous work (18,24–27). In a non-dependent state (pre-CIA), female rats drank more alcohol than their male counterparts. However, CIA males drank more than same-sex controls during acute withdrawal, which is a hallmark of dependence-like behavior (29,30). Although dependence-induced escalation was not observed in females, which is consistent with other studies (18,26), it is important to note that CIA does elicit other signs of dependence in female rats, including increased signs of physical withdrawal and negative affective behavior (18,26,31). We therefore reason that the current data is still relevant to alcohol dependence in females.

There are a few limitations of the current work that require consideration. First, as mentioned above, females exposed to CIA did not show escalated alcohol consumption during acute withdrawal, which is a key component of alcohol dependence (29,46). There are several potential explanations, including that females appear less sensitive than males to the aversive effects of drug withdrawal (47), or that the exposure time of CIA may not have been adequate to induce escalated alcohol self-administration for females. Although as little as ten days of CIA exposure can lead to escalated alcohol intake in male rats (26), other studies have shown escalations in female self-administration only after ten weeks of CIA exposure (27).

This limitation highlights a long-standing issue in preclinical research, namely that there is a dearth of information regarding female behavior and neurobiology compared to males, and that most animal models, including CIA, have been built using only male subjects. The outcomes are often different when these models are retroactively applied to females. This doesn’t necessarily mean that female rats don’t experience alcohol withdrawal or dependence, but that the phenotype may be different in females or that the way we measure behavior is not adequate to capture such phenotypes in female rodents, as has been observed with other behaviors (48,49). Our ongoing and future work aims to tackle some of these issues by determining whether females show escalation after longer periods of CIA exposure, require a different withdrawal period, and/or require more or longer operant sessions.

A second limitation is that we were unable to predict current drinking behavior using LFPs from the tethered self-administration sessions. Tethering animals significantly reduced drinking, to 0 in many cases, despite attempts to habituate animals to the tether. Alcohol drinking behavior, especially during withdrawal, is highly sensitive to stressors that often suppress rather than enhance drinking (50). Our future work aims to rectify this problem by increasing the number of habituation sessions or using wireless recording equipment that would eliminate the need for a tether.

Overall, the current study has allowed us to determine whether and how cortical, striatal, and limbic oscillations are impacted by CIA in males and females, as well as which specific LFP features correlate with drinking. Based on these findings, our future work will determine whether the CIA-induced changes in LFPs observed here are functionally relevant. While LFPs are thought to reflect neuronal activity, the precise nature of the relationship between what is happening at a cellular level and LFPs is poorly understood (51). Our future work will use chemogenetic and optogenetic techniques to selectively target cortical-striatal vs. cortical-limbic circuits in CIA-exposed animals to determine whether manipulation of these circuits will reduce drinking and other measures of dependence in a sex-specific manner, while also ameliorating the effect of CIA on LFPs.

## CONCLUSIONS

Although it is very probable that CIA-induced changes to cortical-striatal and cortical-limbic circuits play a role in both male and female alcohol misuse, the current experiment supports the notion that the neural circuits *most* impacted by chronic alcohol exposure are sex-specific. This study also highlights the vital need to incorporate both sexes into preclinical work, enhancing our ability to translate preclinical findings to clinical populations. For example, DBS targeting the NAc is currently going through clinical trials for treatment-resistant AUD, but previous preclinical and clinical DBS studies were performed almost exclusively in males (52), and targeting different circuits might better benefit women. Understanding that limbic regions might play a stronger role in driving alcohol misuse in females could provide an important biological treatment target for future studies aimed at developing more efficacious therapies for AUD in both sexes.

## DECLARATIONS

### Ethics approval and consent to participate

Animal facilities were accredited by the American Association of Laboratory Animal Care (AALAC), and all procedures were approved by the Institutional Animal Care and Use Committee (IACUC) at Washington State University.

### Consent for publication

Not applicable.

### Availability of data and material

Data sets generated during the current study are available from the corresponding author on reasonable request. Matlab codes to generate the predictive models are available on GitHub: https://github.com/lucasdwi/code/tree/greenlab.

### Authors’ contributions

KH assisted in developing the research design, collected and analyzed the data, and drafted the manuscript. SN and MP assisted in collecting the data and drafting the manuscript. LD assisted in data analysis and drafting the manuscript. AH developed the research question and research design, assisted in analyzing the data, and drafted and edited the manuscript.

## Acknowledgements

Not applicable.

