## Supplemental Material for "Chronic intermittent alcohol yields sex-specific disruptions in cortical-striatal-limbic oscillations"

**Results**

*Active and inactive lever presses*

For male self-administration behavior, a mixed factorial ANOVA revealed a significant effect of session [*F*(4,44) = 4.75, *p*<0.01, *n_p_^2^* = 0.30], a significant effect of group [*F*(1,11) = 8.25, *p*<0.05, *n_p_^2^* = 0.43], and a significant session*group interaction [*F*(4,44) = 3.39, *p*<0.05, *n_p_^2^* = 0.24] for active lever presses. Post-hoc tests showed that CIA exposed males drank more alcohol in sessions 3 and 4 (*p*<0.05) compared to control males. For inactive lever presses, there is a significant effect of session [*F*(4,44) = 5.50, *p*<0.01, *n_p_^2^* = 0.33], but no effect of group [*F*(1,11) = 0.001, *p*=0.97, *n_p_^2^* = 0.00] or a group*session interaction [*F*(4,44) = 0.59, *p*=0.68, *n_p_^2^* = 0.05] (Supplemental Figure 1A).

For female self-administration behavior, a mixed factorial ANOVA revealed a significant effect of session [*F*(4,48) = 3.00, *p*<0.05, *n_p_^2^* = 0.20], but no effect of group [*F*(1,12) = 0.41, *p*=0.54, *n_p_^2^* = 0.03] or a session*group interaction [*F*(4,48) = 0.27, *p*=0.90, *n_p_^2^* = 0.02] for active lever presses. For inactive lever presses, there is a significant effect of session [*F*(4,48) = 3.50, *p*<0.05, *n_p_^2^* = 0.23], but no effect of group [*F*(1,12) = 1.47, *p*=0.25, *n_p_^2^* = 0.11] or a group*session interaction [*F*(4,48) = 1.48, *p*=0.22, *n_p_^2^* = 0.11] (Supplemental Figure 1B).

*Drinking during tethered sessions*

During LFP collection, tethering the animals to the commutator eliminated any group effects on g/kg of alcohol consumed, and drinking was suppressed below baseline levels (Supplementary Figure 2). For both sexes, there was no effect of session [males: *F*(2,22) = 0.004, *p*=0.99, *n_p_^2^* = 0.00; females: *F*(2,24) = 0.22, *p*=0.81, *n_p_^2^* = 0.02], group [males: *F*(1,11) = 0.00, *p*=0.99, *n_p_^2^* = 0.00; females: *F*(1,12) = 0.004, *p*=0.95, *n_p_^2^* = 0.00], or session*group interaction [males: *F*(2,22) = 0.78, *p*=0.47, *n_p_^2^* = 0.07; females: *F*(2,24) = 2.25, *p*=0.13, *n_p_^2^* = 0.16].

*Predicting drinking from LFPs in tethered sessions*

Supplemental Figure 3 shows the average real and permuted model deviance for predicting g/kg consumed during tethered sessions from LFPs. Models in control males (real deviance = 0.25 ± 0.1, permuted deviance = 0.29 ± 0.0; 3A), CIA males (real deviance = 0.69 ± 0.1, permuted deviance = 3.42 ± 2.0; 3B), control females (real deviance = 136.2 ± 103.1, permuted deviance = 2.0 ± 1.3; 3C), and CIA females (real deviance = 0.42 ± 0.05, permuted deviance = 0.44 ± .03; 3D) did not outperform chance estimates.

**Figure captions**

**Supplemental Figure 1:** Average active (AL) and inactive (IL) lever presses in male (A) and female (B) rats exposed to CIA or room air (control) during 30-minute self-administration sessions (pre-CIA baseline) and after (acute withdrawal) four weeks of CIA exposure. CIA led to increased active lever presses in the last two self-administration sessions in males (*p<0.05; n=6-7/group), but no changes in female drinking (n=7/group). There were not significant effects on inactive lever presses.

**Supplemental Figure 2:** Average g/kg of alcohol consumed in male (A) and female (B) rats while recording LFPs. Tethering to the commutator to collect LFPs eliminated the effect of CIA on group in males and significantly reduced rates of self-administration in all groups (n=6-7/group/sex).

**Supplemental Figure 3:** Average model deviance +/- SD for real and permuted continuous prediction models (n = 6-7/sex × 3 recordings/rat) in males (A and B) and females (C and D). Models built to predict g/kg of alcohol consumed during tethered sessions did not outperform chance estimates.

**Figures**

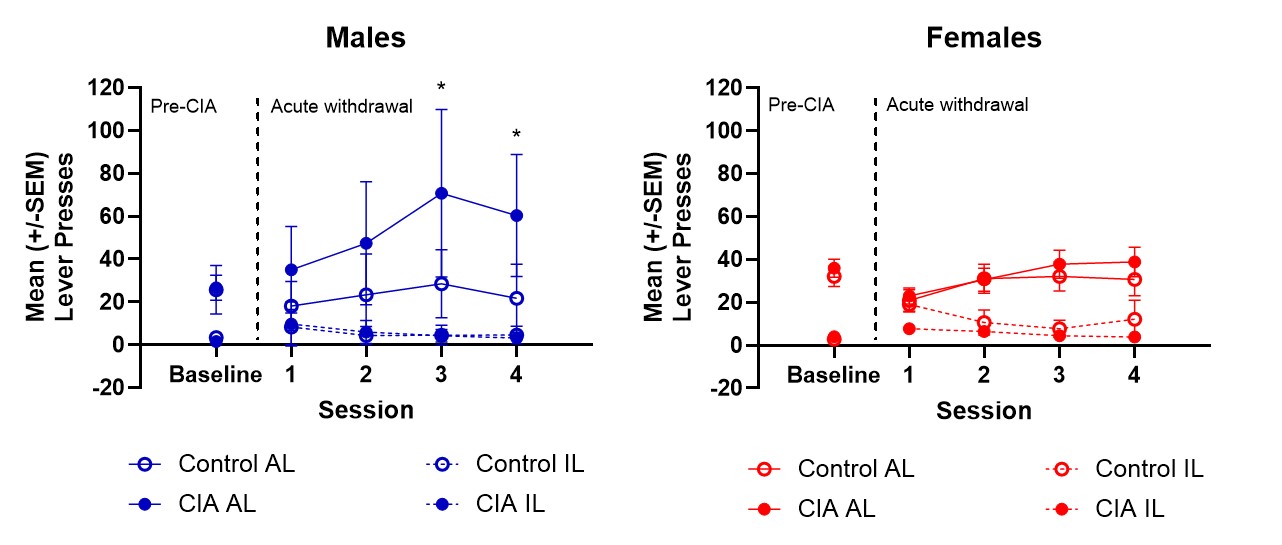

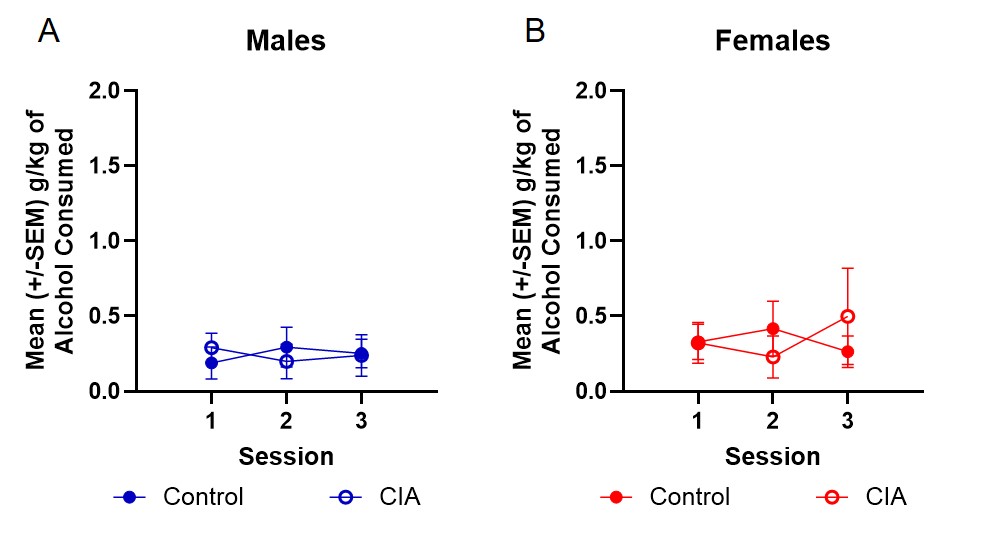

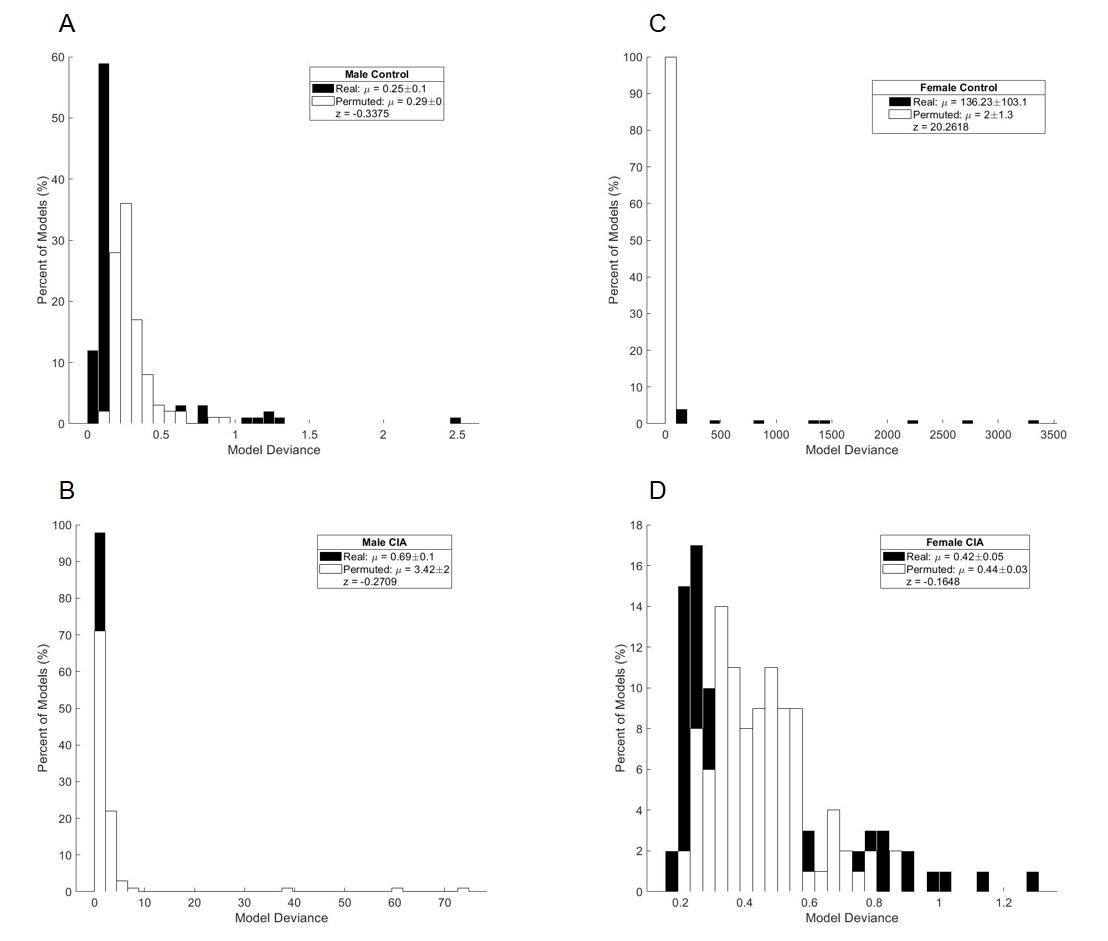
