## Supplementary figures and images for "Chronic intermittent alcohol yields sex-specific disruptions in cortical-striatal-limbic oscillations"

### Supplemental Figure 1

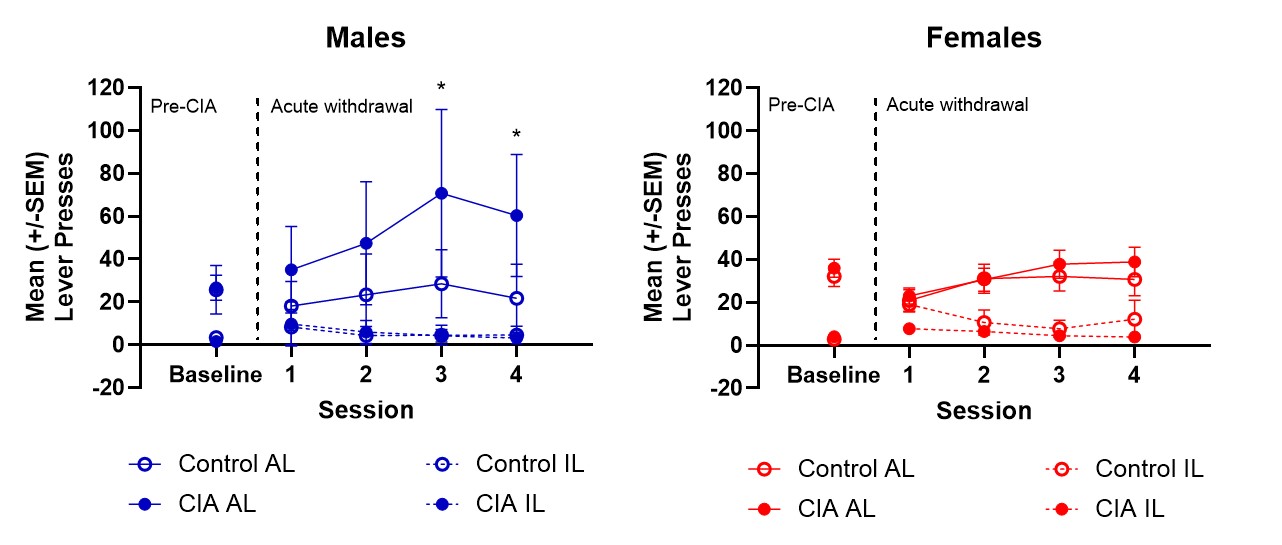

### Supplemental Figure 2

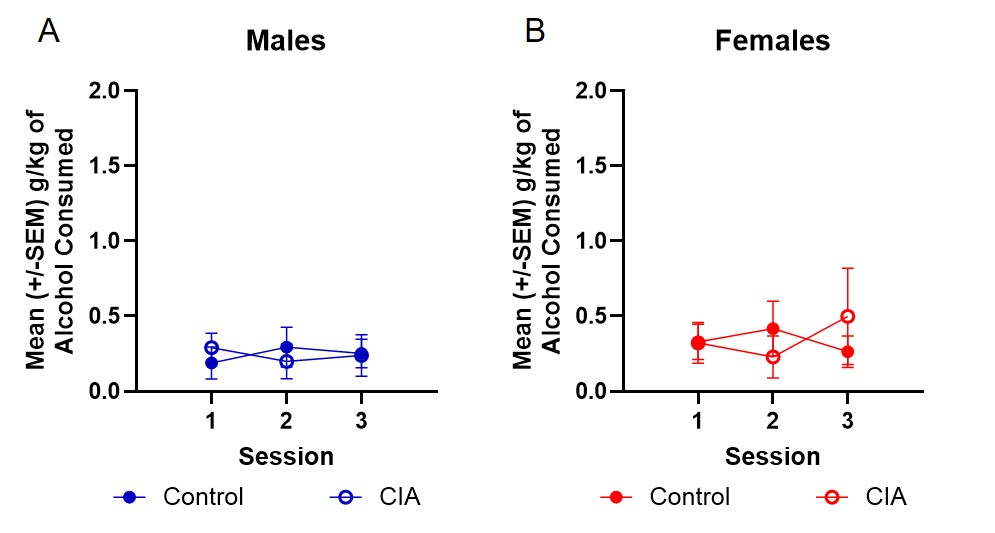

### Supplemental Figure 3

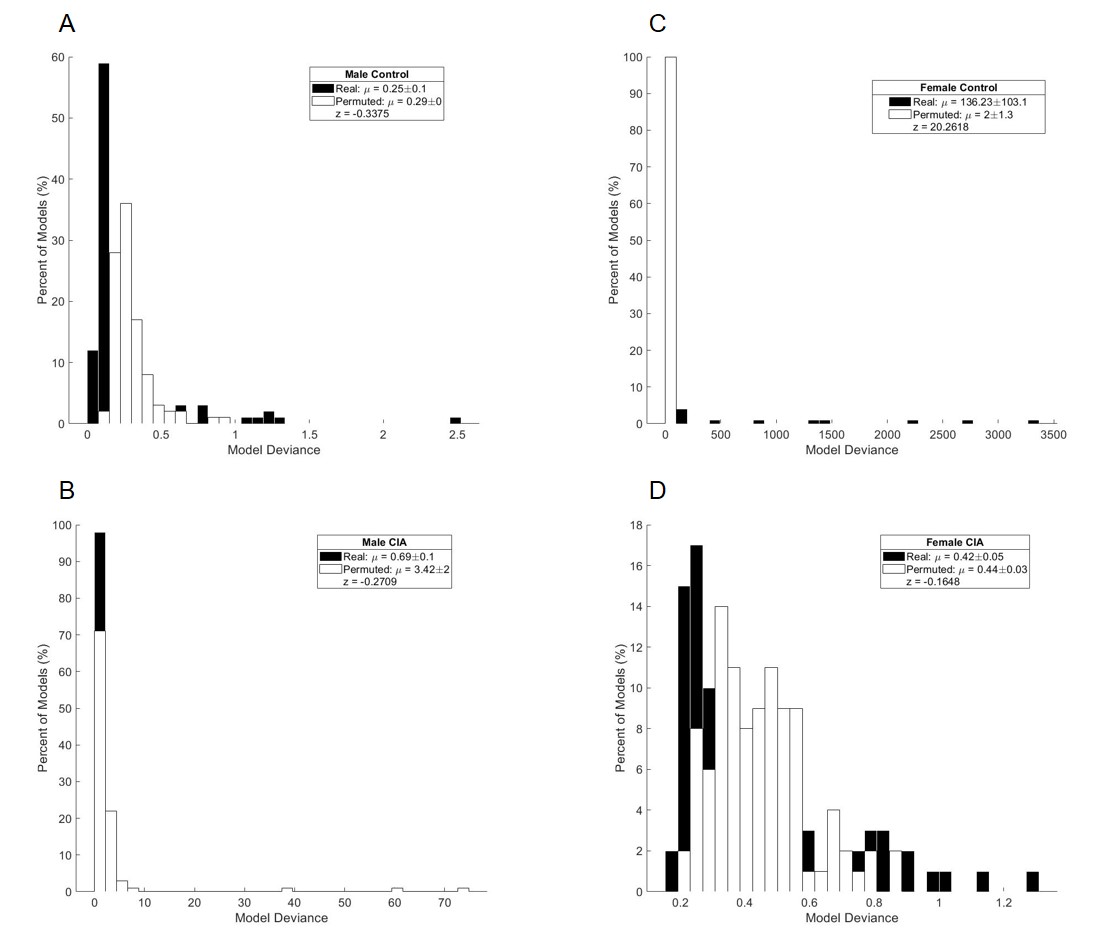
